# Increased elastase sensitivity and decreased intramolecular interactions in the more transmissible SARS-CoV-2 variants’ spike protein: Analysis of the new UK and SA SARS-CoV-2 variants

**DOI:** 10.1101/2021.01.19.427355

**Authors:** S. Pokhrel, L. Lee, B.R. Kraemer, K. Samardzic, D. Mochly-Rosen

## Abstract

Two SARS-CoV-2 variants showing increased transmissibility relative to the Wuhan virus have recently been identified. Although neither variant causes more severe illness or increased risk of death, the faster spread of the virus is a major threat. Using computational tools, we found that the new SARS-CoV-2 variants may acquire an increased transmissibility by increasing the propensity of its spike protein to expose the receptor binding domain. This information leads to the identification of potential treatments to avert the imminent threat of these more transmittable SARS-CoV-2 variants.

**Teaser:** The more infective SARS-CoV-2 variants may expose its Achilles Heel – an opportunity to reduce their spreading.

## MAIN TEXT

### Introduction

Severe acute respiratory syndrome coronavirus 2 (SARS-CoV-2), the novel coronavirus that has resulted in the coronavirus disease 2019 (COVID-19) pandemic, continues to mutate while spreading throughout the world. A critical protein on the surface of the virus is the spike protein, that mediates the entry of the virus into the host cells (*1*). When a mutation or set of mutations provide an advantage over the previous variants, the new variant becomes the dominant variant to spread. This is what occurred when the aspartate 614 in the spike protein of the initial (Wuhan) SARS-CoV-2 virus, mutated to a glycine (Asn614Gly); within a month, the new variant became the most dominant in Europe (*2*) and now globally (*3*). The Asn614Gly variant is associated with higher viral loads, but not with a more severe COVID-19 symptoms in infected individuals (*4*). Importantly, the Asn614Gly mutation does not affect the ability of antibodies to neutralize that variant (*5*). Nevertheless, as the virus continues to mutate at a very high rate, [for example, each amino acid in the 1273 amino acid of the viral spike protein has mutated on average 4 times since the protein was first sequenced, about a year ago (*6*)], the need to continue active surveillance for the emergence of new variants and examination of means to slow down the spread of the new variants is of utmost importance.

### The SARS-CoV-2 United Kingdom variant, 501Y.V1, and the South African variant, 501Y.V2

A novel SARS-CoV-2 variant, known as B.1.1.7 or 20B/501Y.V1, emerged in the United Kingdom at the end of September 2020 and became the dominant variant within a month (*7*). This variant increases viral transmissibility between 40-70% (*8*). As of January 16 2021, 501Y.V1 has been reported in more than 50 countries (*9*). However, this new variant does not appear to increase the severity of COVID-19 (*10*). The variant has concomitant 3 deletions and 10 amino acid changes in the 1273 amino acid spike protein (Table 1a), compared with the initial SARS-CoV-2 index virus identified in Wuhan, China; only one (Asn501Tyr or N501Y) is in the angiotensin-converting enzyme 2 (ACE2) receptor-binding domain (RBD; amino acids 331-524) (*11*). Since this Asn501Tyr mutation in the spike’s RBD was observed alone as early as April 2020 in Brazil and later in Australia without reports of increased transmissibility, (*12*), it is unlikely that this single substitution is sufficient to explain the new phenotype of B.1.1.7 variant.

**Table 1:**
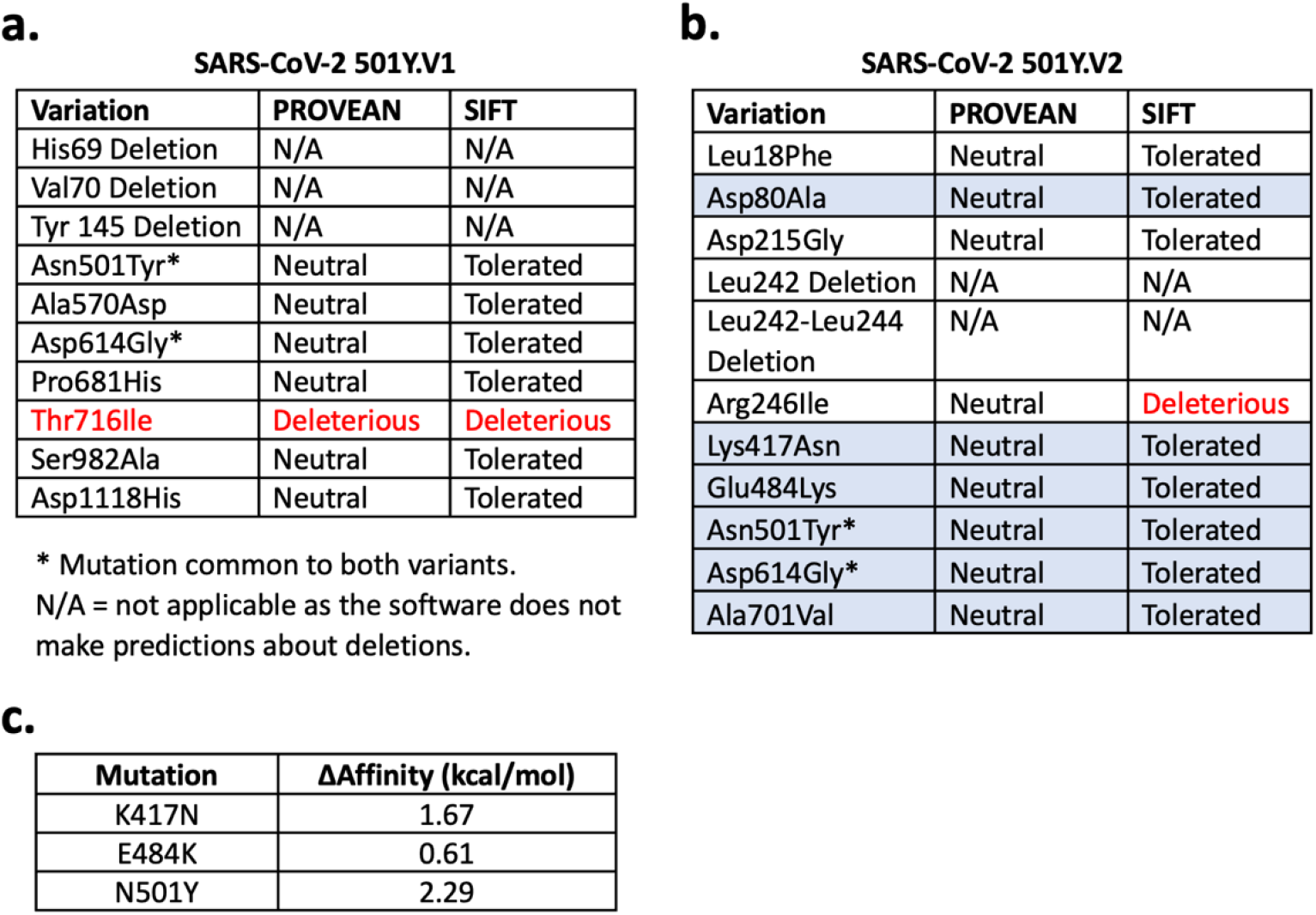
Predicted biological impact of mutations in the spike protein and ΔAffinity calculations. Predicted effects of amino acid substitutions common in **(A)** SARS-CoV-2 501Y.V1 and **(B)** SARS-CoV-2 501Y.V2 using Protein Variation Effect Analyzer (PROVEAN) (*37*) and SIFT (*40*). Invariant (fixed) mutations in all the 501Y.V2 isolates are shaded blue **(C)**: ΔAffinity calculations for RBD variants.

**SARS-CoV-2 501Y.V1**
| Variation | PROVEAN | SIFT |
| --- | --- | --- |
| His69 Deletion | N/A | N/A |
| Val70 Deletion | N/A | N/A |
| Tyr 145 Deletion | N/A | N/A |
| Asn501Tyr* | Neutral | Tolerated |
| Ala570Asp | Neutral | Tolerated |
| Asp614Gly* | Neutral | Tolerated |
| Pro681His | Neutral | Tolerated |
| Thr716Ile | Deleterious | Deleterious |
| Ser982Ala | Neutral | Tolerated |
| Asp1118His | Neutral | Tolerated |

\* Mutation common to both variants.

**SARS-CoV-2 501Y.V2**
| Variation | PROVEAN | SIFT |
| --- | --- | --- |
| Leu18Phe | Neutral | Tolerated |
| Asp80Ala | Neutral | Tolerated |
| Asp215Gly | Neutral | Tolerated |
| Leu242 Deletion | N/A | N/A |
| Leu242-Leu244 Deletion | N/A | N/A |
| Arg246Ile | Neutral | Deleterious |
| Lys417Asn | Neutral | Tolerated |
| Glu484Lys | Neutral | Tolerated |
| Asn501Tyr* | Neutral | Tolerated |
| Asp614Gly* | Neutral | Tolerated |
| Ala701Val | Neutral | Tolerated |

**c.**
| Mutation | ΔAffinity (kcal/mol) |
| --- | --- |
| K417N | 1.67 |
| E484K | 0.61 |
| N501Y | 2.29 |

De Olivera and collaborators identified another more transmittable and dominant variant, termed 501Y.V2 (aka 20H/501Y.V2 or B.1.351) South African variant (*13*). Identified first in the second week of October, this variant became dominant in South Africa within a month. Six fixed substitutions in all the South African variants were identified (Table 1b): Asn501Tyr [identical to the 501Y.V1 UK variant (*11*)] and Pro614Gly [identical to the European dominant variant, identified between March and April of 2020 (*2*)], the Asp80Ala, Lys417Asn, Glu484Lys and Ala701Val. It appears that the combination of these substitutions results in increased infectivity without increasing COVID-19 severity (*13*).

Here we focused on mutations in the SARS-CoV-2 viral surface spike protein to examine how the new variant became more infective. We used computational methods, including Molecular Operating Environment (MOE) analysis (*14*) and software to predict the outcome of substitutions on the protein structure, to examine the features acquired by the new variants that enable them to increase the rate of infection and spreading without increasing the severity of COVID-19, the pathology resulting from the infection. The goal of this study is to determine whether the acquisition of the more infective phenotype (resulting in a more efficient and rapid viral transmission) exposes new vulnerabilities to target these variants with current drugs.

## Results

### Potential acquired features of the SARS-CoV-2 501Y.V1 and SARS-CoV-2 501Y.V2 variants

The SARS-CoV-2 spike protein is a trimer protein on the surface of the virus. This protein mediates the binding to the human ACE2 receptor on the nasal mucosa (*15*). Each monomer in the trimeric spike protein is found in two confirmations: a closed conformation, in which the ACE2 RBD is folded in and is inaccessible, and the open conformation – induced by proteolysis of the spike protein, in which the RBD becomes exposed (*16*). When examining the location of the various mutations associated with increased transmissibility of 501Y.V1 and 501Y.V2, we found that these mutations are not located in a particular domain in the spike protein, nor are they all exposed in the close and or the open conformation (Fig. 1a-c).

**Fig 1:**
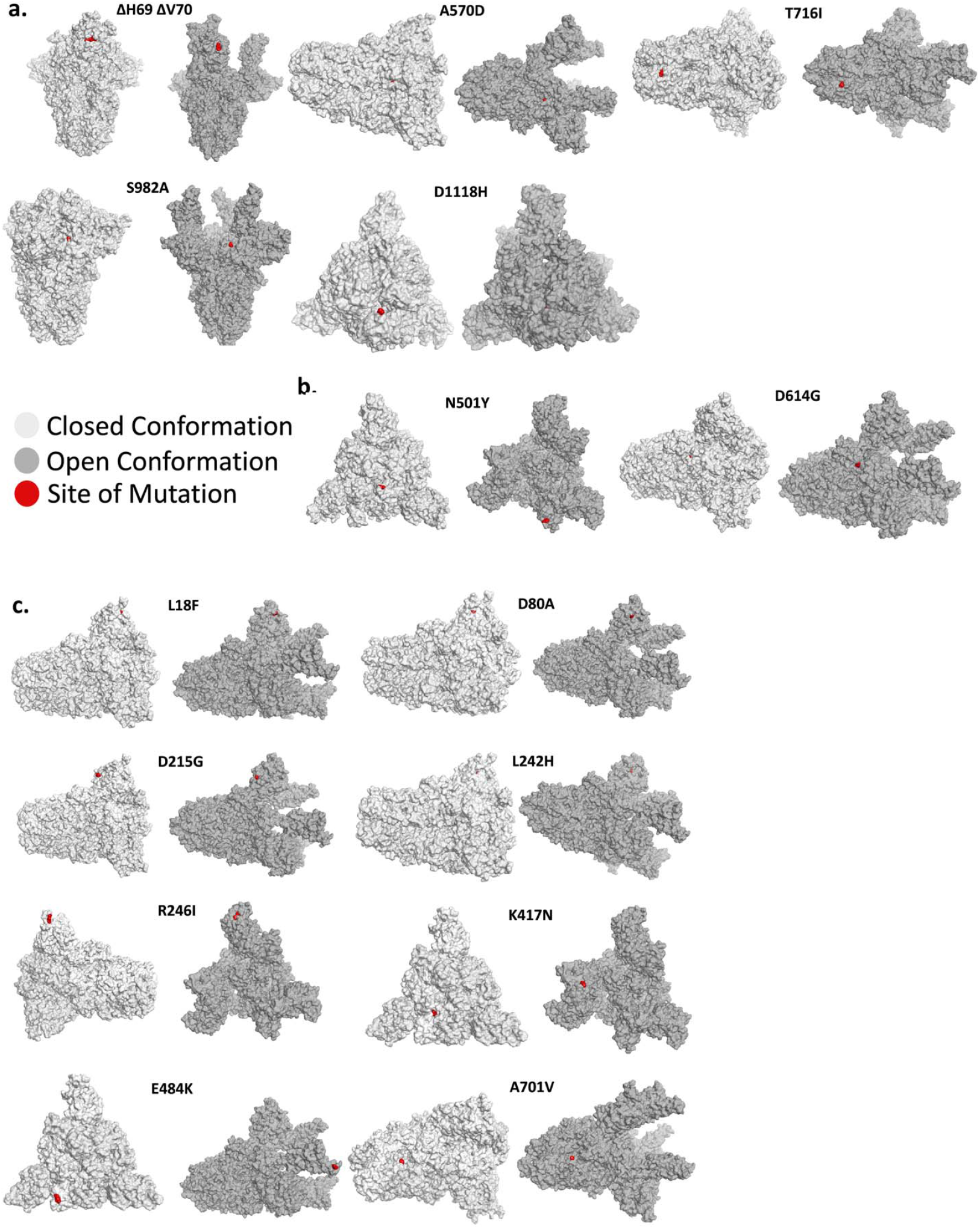
Mutant position in the open and closed conformation of the spike protein. **(A)** The position of various mutations of the Y501.V1, **(B)** The N501Y and D614G mutations, which occur in both the Y501.V1 and Y501.V2 variants, and **(C)** Mutations unique for the 501Y.V2. Each mutation is shown in red in one of the monomers in the 3D structure of the spike protein trimer. Shown is the closed conformation (light gray) and the open (dark gray) conformation.

As neither variant with increased transmissibility causes more severe COVID-19 disease (*3, 10*), and the mutations do not cluster to a particular site in the spike protein, we next examined the possibility that the acquired features in the new variants may lie in increased susceptibility of the spike protein to activation due to introduction of additional protease-activating sites. We hypothesized that the proteases responsible for this gain-of-function should be relatively unique to the initial site of SARS-CoV-2 entry, the nasal mucosa, as viral effects on internal organs do not seem to become more severe. At least seven ubiquitously expressed proteases have been suggested to act on the spike protein: TMPRSS2, furin, PC1, matriptase, trypsin, cathepsin L and/or cathepsin B (*17, 18*).

In contrast to the above proteases, the neutrophil elastase is enriched in the nose of humans compared to other tissues (*19*) and neutrophils, a source for this elastase, are recruited to the nose in SARS-CoV-2 infection (*20*). Indeed, the now worldwide dominant Asp614Gly variant, introduced a new elastase proteolysis site in the spike protein (*2, 21*). Using ExPASy (*22*), we identified Thr716Ile and Ser982Ala in 501Y.V1 as potential new elastase cleavage sites (Fig. 2a). However, whereas Thr716 is on the surface of the spike protein, Ser982 is barely exposed in the open (active) conformation and is buried in the closed conformation (Fig. 1a). We also identified new potential elastase cleavage sites in 501Y.V2: two of the substitutions are fixed in all variants of the new South African strain (Asp80Ala and Ala701Val) and the other two are not (Asp215Gly and Arg246Ile; Fig. 2b). Note that although proteolysis-induced activation was first mapped to the S1/S2 boundary in the spike protein (*23*), protease-induced cuts at other sites may increase the propensity of the spike protein to expose the RBD.

**Fig. 2:**
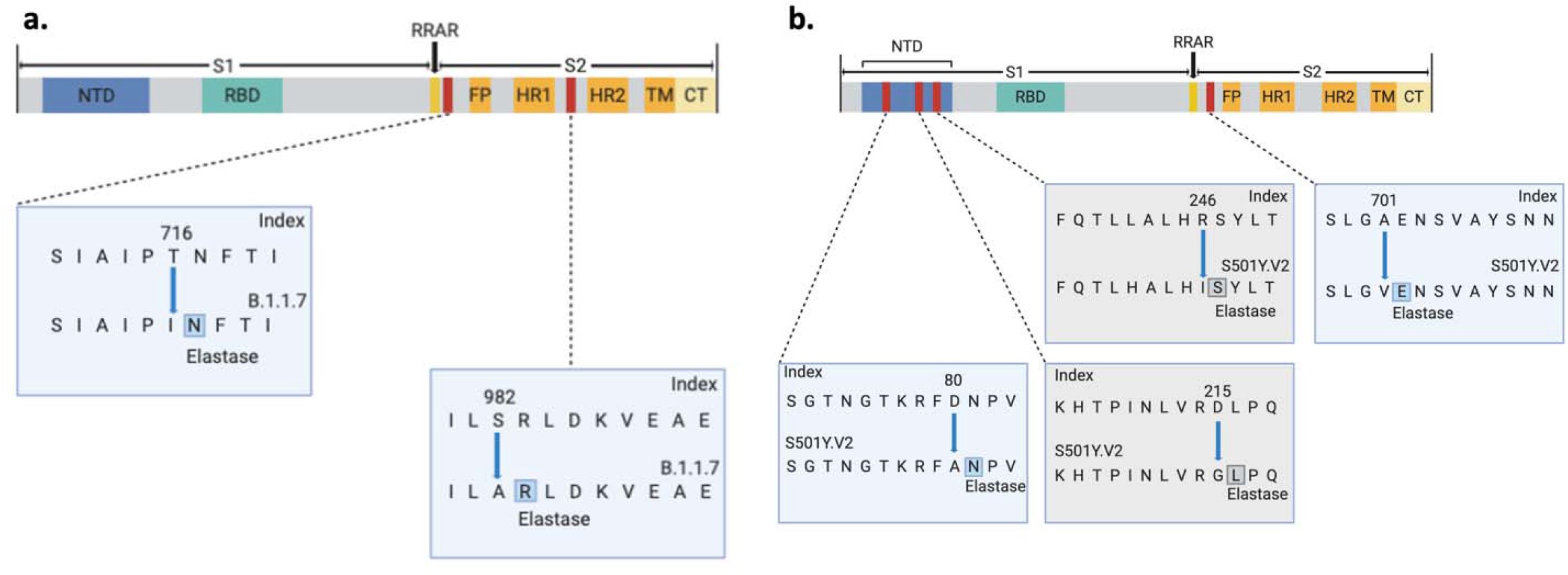
Potential elastase sites in the spike protein of the new SARS-CoV-2 variants. Potential new elastase sites in **(A)** the more infective UK variant, 501Y.V1 and **(B)** the more infective South Africa variant, 501Y.V2. Images generated using BioRender. Blue boxes indicate fixed mutations and gray boxes – non-fixed in all the sequenced 501Y.V2 (*13*).

### The new variants may decrease intramolecular interactions in the spike protein

We next examined a complementary possible mechanism for the new mutations to compete for the older viral variants, namely substitutions that may decrease intramolecular interactions in the spike protein, thus rendering it more likely to be in the open/active conformation. The open confirmation in this trimeric protein exposes the RBD in a monomer, thus enabling it to bind to ACE2 and enter the body. Indeed, previous work suggested that the Asp614Gly substitution is more prone to be in the open conformation (*24*).

Using MOE, we found that Asn501Tyr substitution in both 501Y.V1 in 501Y.V2 may increase the propensity of the spike protein to be in the ‘open’ or active conformation (Fig. 3a). The backbones of Asn501 residues in the three monomers are about 14 Å from each other in the closed trimeric structure. Therefore, a substitution with a bulky tyrosine residue (∼7 Å, each) may introduce clashes and cause a loosening of the contact sites between the monomers in the closed conformation (Fig. 3a). Note, however, MOE calculations suggest that 501Tyr substitution does not significantly destabilize the closed conformation (ΔStability 0.4369 kcal/mol) as compared with the index Wuhan 501N variant.

**Fig. 3:**
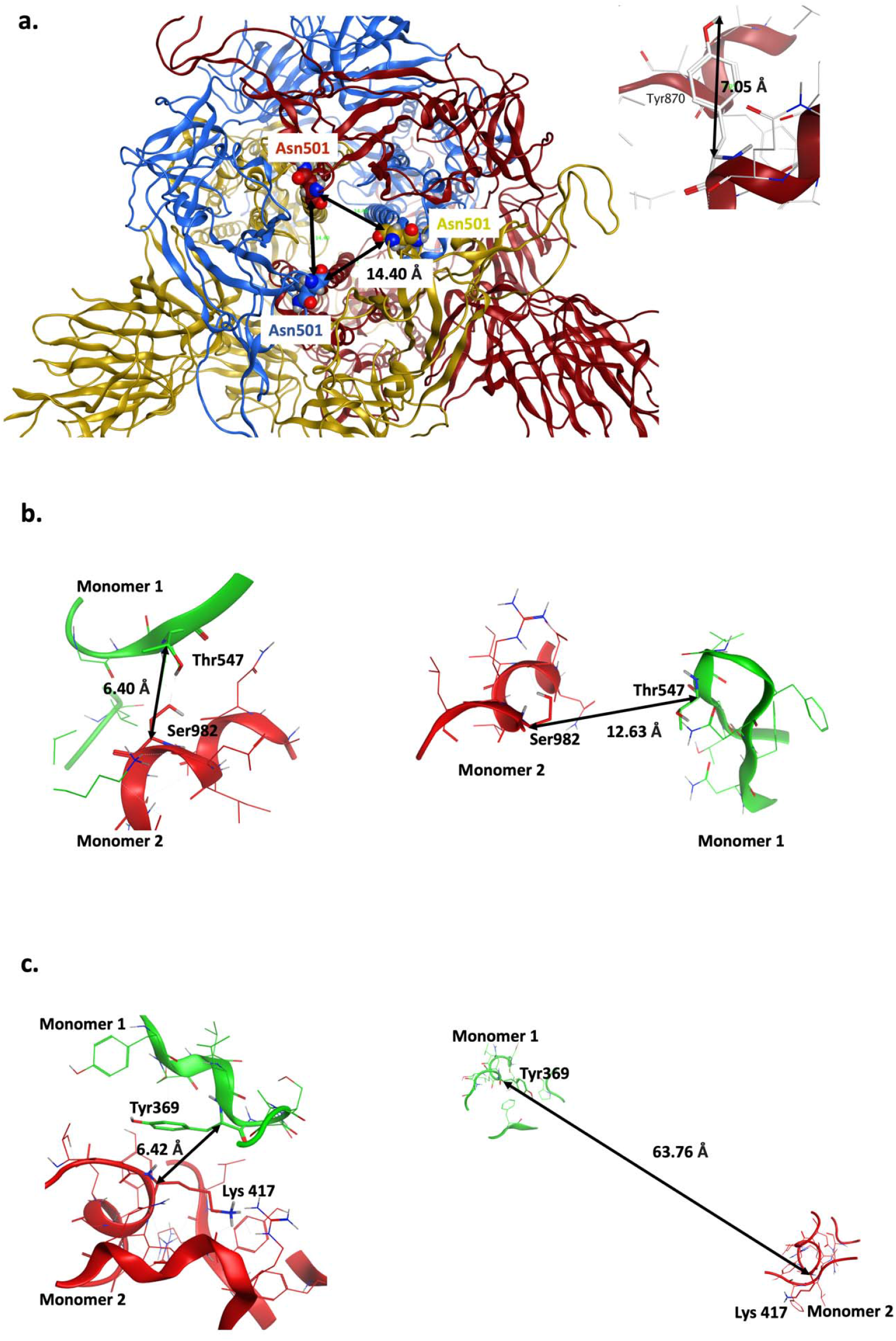
Potential destabilizing effects of the point mutations on the closed conformation of the spike protein trimer. **(A)** Distance between the backbones of Asn501 in closed conformation is shown (left) and typical length of Tyr side chain is shown (right). Asn501Tyr mutation might introduce steric clashes and destabilize the closed conformation. **(B**) Ser982 (monomer 2)-Thr547 (monomer 1) interaction in closed (left) and open (right) conformation. Ser982Ala results in loss of this inter-monomeric interaction. **(C)** Lys417 (monomer 2)-Tyr369 (monomer 1) interaction in closed (left) and open (right) conformation. Lys417Asn mutation may reduce this inter-monomeric interaction.

Another intramolecular interaction in 501Y.V1 that can affect transition from closed to open conformer is mediated by serine 982. Ser982 in one monomer forms a hydrogen bond with a Thr547 in an adjacent monomer in the closed conformation (Fig. 3b). The substitution serine to alanine in 501Y.V1 abolishes this interaction with Thr547 in the neighboring spike monomer, thus likely favoring the open conformation.

In addition to the Asn501Tyr, the South African variant, 501Y.V2, has another amino acid substitution that may favor open conformation of spike protein: Lys417Asn (Table 1b). In the closed conformation, Lys417 present in one monomer makes a proton-π interaction with Tyr369 present in the adjacent monomer (Fig. 3c). Asparagine substitution at this position may disrupt this interaction with Tyr369, potentially favoring the transition to open conformation.

Note also that Thr716Ile, Ser982Ala, and Asp1118His are also conserved in viruses isolated from bat, civet and pangolin, suggesting that these amino acid substitutions may be important for some function of the spike protein (Fig. 4). Similarly, Asp80Ala and Ala701Val amino acid that are mutated in 501Y.V2 are conserved in these other species (Fig. 4). Whether these mutations are associated with more efficient transmissibility in various species remain to be determined.

**Fig. 4:**
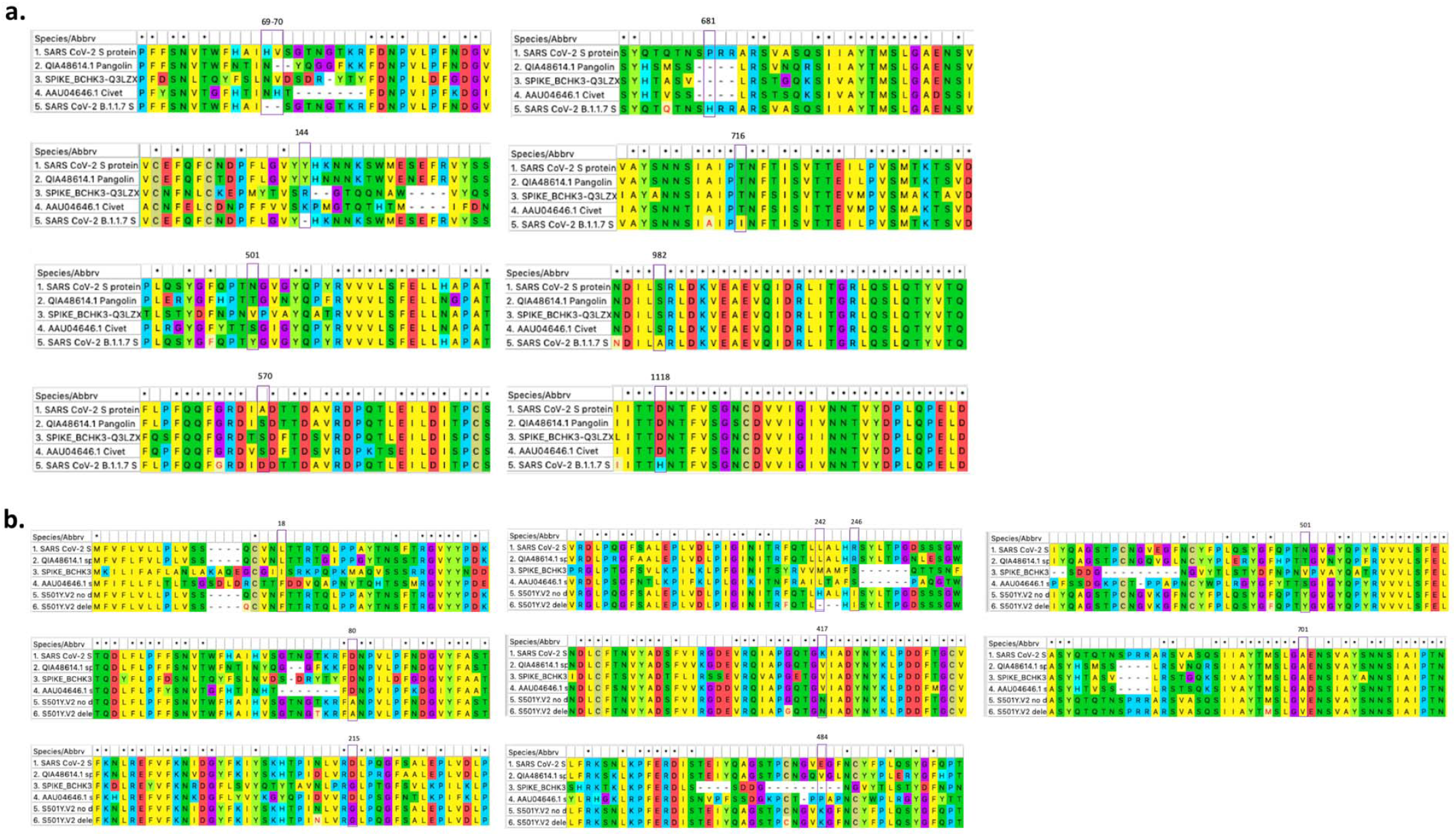
Sequence alignment of the spike protein of the SARS-CoV-2 variants. **(A)** Conservation of mutations found in 501Y.V1 in comparison to the Wuhan index human, bat, civet and pangolin spike protein sequences. **(B)** Conservation of mutations found in 501Y.V2 in comparison to the index Wuhan human, bat, civet and pangolin spike protein sequences. Sequence alignments generated using MEGA X software.

### SARS-CoV-2 variants in the RBD

The Ala501Tyr mutation common to both 501Y.V1 and 501Y.V2 variants and Glu484Lys and Lys417Asn of the 501Y.V2 variant is among the amino acids shown to interact with ACE2 (*25*). It is therefore possible that these mutations increase the binding of the active conformer of the spike protein to ACE2 or to other associated receptors to mediate more effective transmissibility. Although the affinity calculations in MOE suggest that these mutations have minimal negative impact on the affinity of RBD with ACE2 (ΔAffinity; Table 1c), recent yeast display assay suggests that co-occurrence of these mutations increases the affinity of RBD to ACE2 by 2 and 10 folds for 501Y.V1 and 501Y.V2, respectively (*26*). However, we suggest that mutations in RBD may also affect the closed conformation’s stability, thus increasing the probability of binding to ACE2; the monomers wrap around one another through RBD in the closed trimeric structure and mutations in RBD that abolish these critical inter-monomeric interactions could favor the transition to open conformation. Therefore, the effect of these mutations, alone or in combination, on RBD-ACE2 affinity should be tested using intact viral particles.

## Discussion

### Mutations and the impact on SARS-CoV-2 neutralizing antibodies

The emergence of new virus variants triggers the concern that previous exposure to the virus, antibody therapeutics and current vaccines against SARS-CoV-2 may not be effective. This public health concern needs to be addressed as soon as possible. Many neutralizing antibodies were mapped to the RBD, and four major classes of neutralizing sites have been identified in this domain (*27, 28*). As one of the mutations in SARS-CoV-2 501Y.V1 and three mutations in 501Y.V2 are in the RBD, these mutations may decrease the neutralizing activities of antibodies (*13*). However, neutralization studies refute this idea; both sera of convalescent patients and immunized subjects appear effective (*29-31*). Notably, mutation E484K in the 501Y.V2 lineage reduces serum binding and neutralization in circulating SARS-CoV-2 isolates (*32*). Furthermore, although point mutations in the spike protein may affect the therapeutic benefit of one or a combination of two monoclonal antibodies (*33*), the benefit of polyclonal antibodies, such as those generated in infected individuals or when using either passive or active vaccines is less likely to be negatively impacted.

### The increased transmissibility of the new variants may expose an Achilles Heel

Our analysis indicates that many of these mutations may also increase the exposure of the RBD of the spike protein. We therefore suggest that neutralization by antibodies may increase – more antigenic determinants per each viral particle may be exposed for the antibodies to bind to. Indeed, sera from hamsters infected with Asp614Gly virus exhibit slightly higher neutralization titers against the new dominant Gly614 variant than against the index variant Asp614 (*13*). Furthermore, each viral particle has as many as 80 spike proteins on each virus (*34*), only some of which need to bind ACE2, to induce viral entry into cells. If more spike proteins are activated and expose the antigenic RBD domain, neutralizing antibodies have an increased probability to bind to viral particles and agglutinate several particles together. The resulting agglutinated particles are less likely to have a successful fusion with the cell membrane and infection may be aborted. Increased infection efficiency due to increased propensity of the spike protein activation may also increase the vulnerability of the usually hidden spike protein core to the neutralizing antibodies.

### Nasal neutrophil elastase and viral infections – a therapeutic opportunityΔ

Proteomic analyses identified five proteins with increased expression in nasopharyngeal swab of subjects with high polymerase chain reaction titer for SARS-CoV-2 (*20*). Among these five proteins is neutrophil elastase, whose levels increased 3-fold in SARS-CoV-2 infected *vs.* noninfected individuals. Furthermore, there is a 10-fold increase in neutrophil number in swabs from infected individuals. Elastase was also found to be critical for accelerating viral entry as well as inducing hypertension, thrombosis and vasculitis (*20, 35*). As there are several approved elastase inhibitors, including Sivelestat (which is approved for acute respiratory syndrome), Alvestat (α1-antitrypsin) and Roseltide, the use of these inhibitors, perhaps as intranasal spray or drops, should be considered.

### Limitations to our analysis

Our study uses computational tools to examine the potential impact of the mutations found in SARS-CoV-2 501Y.V1 and 501Y.V2 on the viral infectivity has several limitations. We focused only on two aspects – sensitivity to protease and increased open conformer of the spike protein. Other mechanisms may increase viral transmissibility and were not examined here. For example, if proteolytic events contribute to viral inactivation at the mucosa, a decline in proteolysis susceptibility, especially to proteases present in the nose, may also contribute to increased infectivity. We also did not examine the possibility that the increased transmissibility of the new variants may be due to better survival of the virus in respiratory droplets, the potential increased affinity of the viral spike protein for receptors other than ACE2, nor the impact of mutations in other viral genes. Finally, our analysis tested mainly the potential impact of one mutation at a time. As our analysis is *in silico*, experiments to test the hypotheses raised by this study should be conducted next, to determine if these SARS-CoV-2 variants with more efficient and rapid transmission capabilities are also more sensitive to existing therapies and suggest new therapeutic targets to slow down and arrest the COVID-19 pandemic.

## Materials and Methods

### Proteolysis Site Prediction Using ExPASy

The amino acid sequence of spike protein from the reference EPI_ISL_402124 (Wuhan) sequence was uploaded to ExPASy (*22*) (https://web.expasy.org/peptide_cutter/) and neutrophil elastase was selected as the protease. We examined also potential proteolysis site for the seven other proteases suggested to cleave SARS-CoV-2 spike proteins (*17*). No changes in cleavage sites for both 501Y.V1 and 501Y.V2 were observed for furin, Cathepsin L/B, and PC1. However, a loss of a cleavage site was observed for matriptase in the 501Y.V1, due to alanine of serine 982 mutation. For TMPRSS2 and trypsin, which cleave at single arginine or lysine residues, a proteolysis site was gained at position 484, and a loss of proteolysis site was observed at both positions 246 and 417 in the 501Y.V2.

### Protein Structures

Molecular Operating Environment (MOE) software (version 2019.01) (*14*) was used to generate representative structural models of the open and closed conformations of SARS-CoV-2 variant spike proteins. Protein Data Bank (PDB) ID: 7A98 (*25*) was used to model open conformation whereas PDB ID: 6ZB5(*36*) was used for closed conformation.

### Residue Scan and Affinity Calculation

PDB ID: 7A98 was prepared using the QuickPrep functionality at the default settings, to optimize the H-bond network and perform energy minimization on the system in MOE. The ΔAffinity calculations were performed using 7A98.A (S-protein monomer) and 7A98.D (ACE-2) chains with 7A98.D defined as ligand. Residues K417, E484 and N501 in S-protein (7A98.A) were selected and residue scan application was performed. The difference in affinity (ΔAffinity (kcal/mol)) between mutated residue and the wild-type were calculated as per MOE’s definition.

### Residue Scan and Stability Calculation

QuickPrep functionality was used as above to prepare PDB ID: 6ZB5 in MOE. 6ZB5.A, 6ZB5.B and 6ZB5.C chains were used to perform the calculations. N501 residue in S-protein in all three chains were selected and residue scan application was performed for the observed variants S, Y, T and R that are tolerated at this position with site-limit set to 3. The change in stability (ΔStability; kcal/mol) between the variants and the reference wild-type sequence were calculated as per MOE’s definition.

### Calculating Predicted Effect of Variants in PROVEAN and SIFT

The amino acid sequence of spike protein from the index EPI_ISL_402124 (Wuhan) sequence was uploaded to PROVEAN (*37*) (http://provean.jcvi.org/index.php) and SIFT (*38*) (https://sift.bii.a-star.edu.sg). Every variant observed in 501Y.V1 (UK variant) and 501Y.V2 (South Africa variant) was also uploaded to compare against the reference sequence. Each variant was either predicted to be ‘deleterious’ or ‘neutral’ in PROVEAN or ‘deleterious’ or ‘benign’ in SIFT.

### Multiple Sequence Alignment

MEGA X software (*39*) and the ClustalW alignment function was used to align the amino acid sequences of the human index, bat, civet, pangolin, 501Y.V1 and 501Y.V2 SARS-CoV-2 spike proteins.

## Abbreviations

ACE2: Angiotensin converting enzyme 2
MOE: Molecular Operating Environment
PDB: Protein Data Bank
RBD: Receptor binding domain

## General

We are grateful to the many investigators throughout the world that continue to provide SARS-CoV-2 sequences to public databases.

## Funding

Supported in part by the 2020 COVID-19 Response: Drug and Vaccine Prototyping Grant from the Innovative Medicines Accelerator, Stanford University to D. M.-R. and by the SPARK at Stanford community.

## Author contributions

S.P, L.L and B.R.K provided data analysis, visualization and draft writing. K.S contributed to visualization and manuscript preparation. D.M-R conceived the project and supervised the analysis and writing.

## Competing interests

The authors have no competing interests to declare.

## Notes

### Competing Interest Statement

The authors have declared no competing interest.

